# Activated macrophages restrict invasive bacterial infection in a human intestinal organoid co-culture model

**DOI:** 10.64898/2026.09.02.748782

**Authors:** Lisa F. Goertz, Marcella Cipelli, Manuela Buettner, Guntram A. Grassl, Matthias Lochner

**Affiliations:** Institute of Medical Microbiology and Hospital Epidemiology, Hannover Medical School, Hannover, Germany; Department of Immunology, Institute of Biomedical Sciences, University of São Paulo, São Paulo, Brazil; Central Animal Facility, Hannover Medical School, Hannover, Germany

**Keywords:** intestinal organoids, macrophages, host-pathogen interactions

## Abstract

Intestinal organoids provide physiologically relevant models of the epithelial barrier, but lack the immune compartment that critically shapes host responses to infection. Here, we established a human colon organoid-derived monolayer co-culture system with macrophage-like THP-1 cells positioned directly beneath the epithelial layer. The model enabled controlled apical infection with *Listeria monocytogenes* and *Salmonella* Typhimurium while preserving epithelial barrier integrity. PMA-differentiated THP-1 cells reduced intracellular *L. monocytogenes* burden, whereas additional activation with IFN-γ and LPS resulted in a pronounced reduction of both *L. monocytogenes* and *S.* Typhimurium, accompanied by decreased infection-associated cytotoxicity. Bulk RNA sequencing revealed a distinct co-culture transcriptional signature characterized by coordinated changes in inflammatory, antimicrobial, and epithelial lineage-associated programs. These included reduced HLA-D/MHC class II-associated gene expression, altered *S100A8/S100A9* expression, and changes in epithelial lineage markers indicating a shift in epithelial cellular composition and differentiation state. Together, these findings establish a versatile human organoid-macrophage platform for dissecting epithelial-immune interactions and macrophage-associated control of invasive bacterial infection.

## Introduction

Since the development of organoid technology, substantial progress has been made in refining in vitro models of the gastrointestinal tract. By maintaining a three-dimensional, polarized architecture with physiologically relevant cell–cell interactions, gastrointestinal organoids recapitulate key structural and functional properties of their tissue of origin (1–3). This provides important advantages over conventional planar cancer-derived cell lines, which often display altered signaling pathways and lack the cellular diversity of the intestinal epithelium (4; 5).

Human-derived organoids additionally facilitate the translation of experimental findings to human physiology (6). This is particularly relevant for the study of pathogens that exhibit species-specific host interactions. For example, investigation of *Listeria monocytogenes* infection in conventional mouse models is complicated by species-specific differences in bacterial entry receptors. Although transgenic mice expressing human E-cadherin provide one experimental approach to overcome this limitation (7), human intestinal organoids offer a complementary and physiologically relevant system in which host–pathogen interactions can be studied directly in human epithelial cells. At the same time, organoid-based approaches contribute to the principles of Replacement, Reduction, and Refinement (3Rs) by providing experimental systems that can reduce the reliance on animal models (8; 9).

However, conventional three-dimensional intestinal organoids also present technical limitations for infection studies. Their enclosed architecture restricts experimental access to the apical surface, thereby complicating exposure to pathogens that naturally encounter the intestinal epithelium from the luminal side. Organoid-derived epithelial monolayers grown on permeable supports overcome this limitation by providing independent access to the apical and basolateral compartments while retaining key properties of the intestinal epithelial barrier (10; 11).

Although organoid monolayers provide an advanced model of the intestinal epithelium, the intestinal mucosa comprises multiple additional cell populations that influence epithelial homeostasis and responses to infection (12; 13). In particular, macrophages represent a major immune cell population within the intestinal lamina propria and are positioned in close proximity to the epithelial barrier. Intestinal macrophages contribute to tissue homeostasis and remodeling, surveillance of luminal signals, and the coordination of immune responses during infection (14–16).

Epithelial-immune cell co-culture systems have been established using intestinal organoids and different immune cell populations (17–19). However, their application to defined bacterial infection models remains comparatively limited. In particular, the contribution of basolateral macrophages to epithelial responses during infection with invasive intestinal pathogens remains incompletely understood.

Here, we established a human intestinal organoid-derived monolayer co-culture system in which macrophages are positioned directly beneath the epithelial layer on the basolateral side of a porous Transwell membrane. Using *L. monocytogenes* and *Salmonella enterica* serovar Typhimurium (*S.* Typhimurium) as model pathogens, we investigated how the presence and activation state of macrophages influence bacterial infection and epithelial responses. This platform provides a controlled experimental system for studying epithelial–immune cell interactions during intestinal infection.

## Results

### Establishment and characterization of human colon organoid monolayers

To provide direct experimental access to the apical surface of the intestinal epithelium, we established an organoid-derived monolayer system from human colon biopsies. Human colonic organoids were dissociated into single cells and seeded onto 1-µm-pore Transwell membranes. Cells were expanded for six days to establish confluent epithelial monolayers, followed by an additional two days in differentiation medium (Figure 1A and B). Immunofluorescence microscopy of undifferentiated monolayers at day 6 and differentiated monolayers at day 8 demonstrated the formation of a confluent epithelial layer (Figure 1C). Zonula occludens-1 (ZO-1) staining revealed continuous intercellular junctions throughout the monolayer, whereas apically enriched F-actin staining in cross-sectional images indicated epithelial polarization. UEA-I staining further demonstrated the presence of mucus-associated structures, which were more prominent following differentiation (Figure 1C). To characterize changes in epithelial differentiation, expression of lineage-associated marker genes was analyzed by RT-qPCR (Figure 1D). Differentiation resulted in a significant reduction in expression of the intestinal stem cell marker *LGR5*, accompanied by increased expression of the enterocyte marker *ALPI* and the goblet cell marker *MUC2*. Expression of the secretory progenitor marker *ATOH1* did not change significantly. The development of epithelial barrier function was monitored by measuring transepithelial electrical resistance (TEER). TEER progressively increased during monolayer formation between days 2 and 8 (Figure 1E). To determine the effect of differentiation on barrier properties, TEER values at day 6 were compared with values obtained after an additional 48 hours in either monolayer or differentiation medium. A significant further increase in TEER was observed following differentiation, whereas continued culture in monolayer medium did not result in a significant increase (Figure 1F). Together, these results demonstrate the formation of differentiated and polarized human colonic epithelial monolayers with an intact barrier suitable for subsequent co-culture and infection experiments.

**Figure 1.**
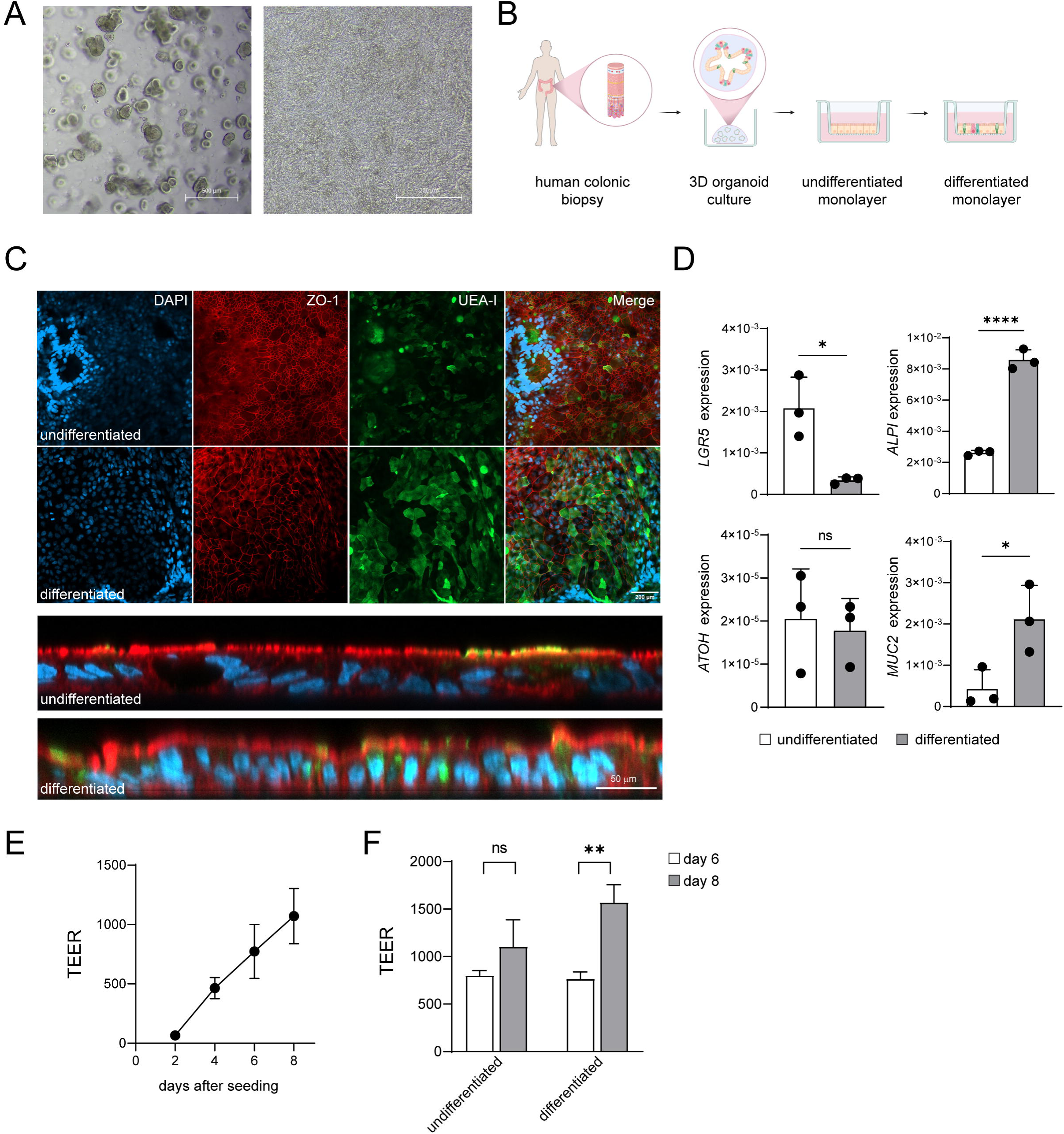
Establishment and characterization of human colon organoid monolayers. (**A**) Representative brightfield images of human colon organoids seven days after seeding (left) and a differentiated organoid-derived monolayer at day 8 (right). Scale bars, 500 µm (left) and 200 µm (right). (**B**) Schematic overview of the generation and differentiation of human colon organoid-derived monolayers on Transwell inserts. (**C**) Representative top-view (top) and cross-sectional (bottom) fluorescence images of undifferentiated and differentiated human colon organoid monolayers. Top view: nuclei (DAPI, blue), ZO-1 (red), and UEA-I (green). Cross-section: nuclei (DAPI, blue), F-actin (phalloidin, red), and UEA-I (green). Scale bars, 200 µm (top) and 50 µm (bottom). (**D**) Relative expression (2^-ΔCt^) of *LGR5*, *ALPI*, *MUC2*, and *ATOH1* in undifferentiated and differentiated human colon organoid monolayers determined by RT-qPCR. Data are presented as mean ± SD. Two-tailed unpaired Student’s t-test. Representative of two independent experiments with three replicates per group. (**E**) TEER values measured at days 2, 4, 6, and 8 after seeding. Data are presented as mean ± SD. One-way ANOVA followed by Tukey’s multiple-comparisons test. Representative of ≥3 independent experiments with three replicates per group. (**F**) TEER values at day 6 and after an additional 48 h in monolayer or differentiation medium at day 8. Data are presented as mean ± SD. One-way ANOVA followed by Tukey’s multiple-comparisons test. Representative of ≥3 independent experiments with three replicates per group.

### Establishment of an intestinal organoid–macrophage co-culture

To incorporate an immune component into the intestinal epithelial model, we established a co-culture system combining human colon organoid monolayers with macrophage-like THP-1 cells. Following phorbol 12-myristate 13-acetate (PMA)-induced differentiation, THP-1 cells were seeded directly onto the basolateral surface of the Transwell membrane. For this purpose, inserts containing confluent epithelial monolayers were temporarily inverted, allowing the THP-1 cells to attach to the underside of the membrane. Following attachment, the inserts were returned to their original orientation and epithelial differentiation was initiated for an additional 48 hours (Figure 2A). Fluorescence microscopy confirmed that THP-1 cells remained attached to the basolateral surface throughout the co-culture period (Figure 2B). Cross-sectional imaging demonstrated the spatial organization of the model, with the epithelial monolayer and THP-1 cells positioned on opposite sides of the porous Transwell membrane and therefore in close proximity to each other. We next assessed whether the presence of THP-1 cells affected the epithelial phenotype. Expression of *LGR5*, *ALPI*, and *MUC2* was not significantly altered by co-culture, whereas ATOH1 expression was significantly reduced in the presence of THP-1 cells (Figure 2C). Thus, co-culture resulted in only limited changes in the epithelial lineage-marker profile under these conditions.

**Figure 2.**
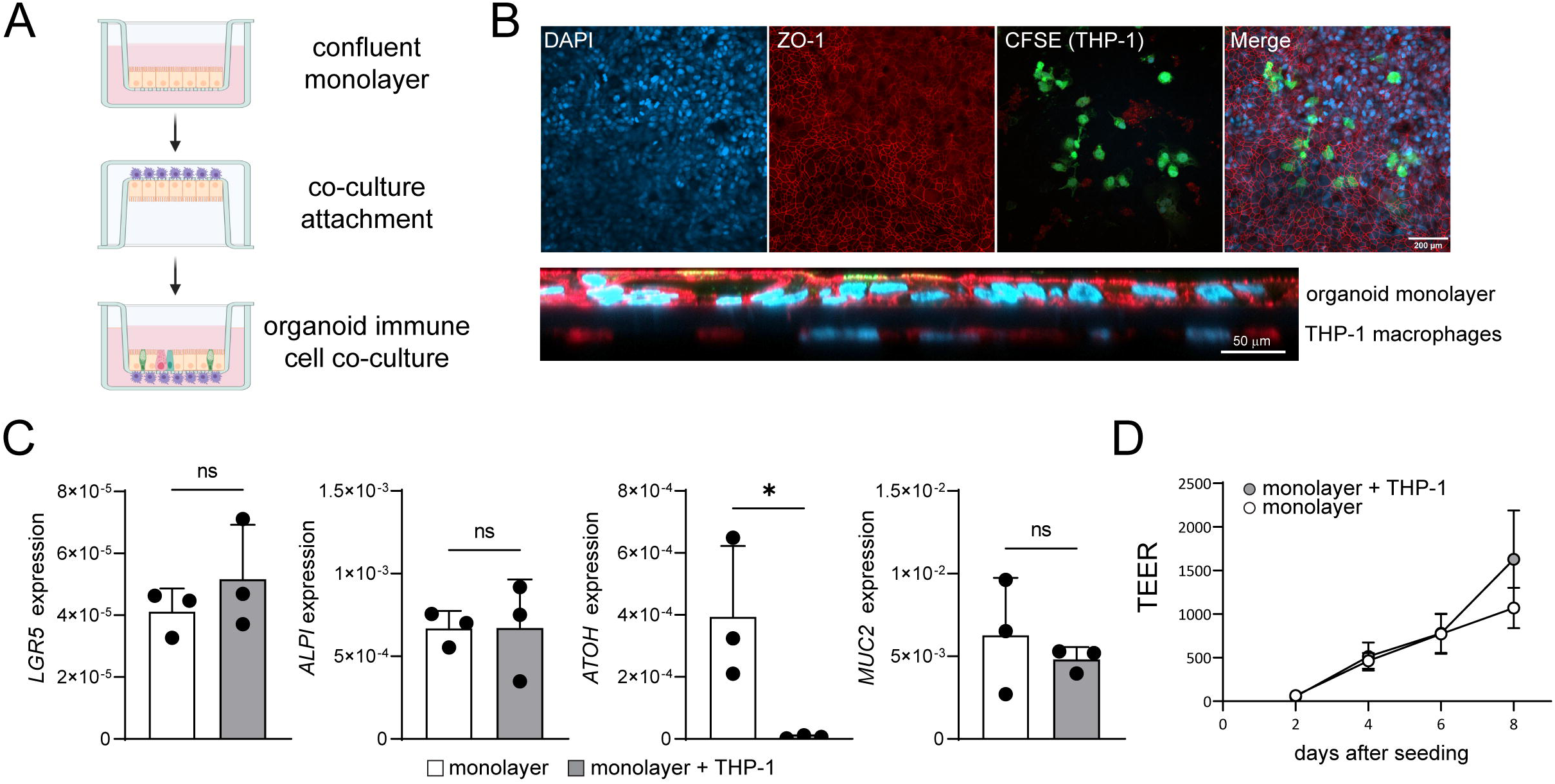
Establishment of the intestinal organoid–macrophage co-culture. (**A**) Schematic overview of the co-culture setup. Confluent organoid-derived monolayers were inverted at day 6, and THP-1 cells were applied to the basolateral side of the Transwell membrane. After 4 h of attachment, inserts were returned to their original orientation. (**B**) Representative top-view (top) and cross-sectional (bottom) fluorescence images of differentiated human colon organoid monolayers co-cultured with THP-1 macrophages. Top view: nuclei (DAPI, blue), ZO-1 (red), and CFSE-labelled THP-1 cells (green). Cross-section: nuclei (DAPI, blue), F-actin (phalloidin, red), and UEA-I (green). Scale bars, 200 µm (top) and 50 µm (bottom). (**C**) Relative expression (2^-ΔCt^) of *LGR5*, *ALPI*, *MUC2*, and *ATOH1* in differentiated human colon organoid monolayers cultured with or without THP-1 macrophages, determined by RT-qPCR. Data are presented as mean ± SD. Two-tailed unpaired Student’s t-test. Representative of two independent experiments with three replicates per group. (**D**) TEER values of human colon organoid monolayers cultured with or without THP-1 macrophages, measured at days 2, 4, 6, and 8 after seeding. Data are presented as mean ± SD. Representative of ≥3 independent experiments with three replicates per group.

Importantly, incorporation of THP-1 cells did not compromise epithelial barrier function. TEER increased progressively during monolayer formation and remained comparable between epithelial monocultures and co-cultures throughout the experimental period (Figure 2D). These findings demonstrate that macrophage-like cells can be stably positioned directly beneath the organoid-derived epithelial monolayer without disrupting overall epithelial barrier integrity, thereby establishing a platform for investigating epithelial–immune cell interactions during infection.

### Activated macrophages restrict bacterial infection in intestinal organoid monolayers

To establish the infection model, differentiated colon organoid monolayers were challenged with the bacterial pathogens *L. monocytogenes* or *S.* Typhimurium. The intracellular bacterial burden was quantified using a gentamicin protection assay and subsequent enumeration of colony-forming units (CFUs) (Figure 3A). Initially, THP-1 cells were differentiated into macrophage-like cells using PMA for 48 to 72 hours prior to co-culture setup. Under these conditions, the presence of THP-1 cells significantly reduced the intracellular burden of *L. monocytogenes*, whereas no significant difference was observed for <u>S.</u> Typhimurium compared with epithelial monolayers alone (Figure 3B). Thus, PMA-differentiated THP-1 cells provided a detectable but pathogen-dependent level of protection under these conditions. To enhance macrophage activation, co-cultures and matched epithelial-only controls were treated with interferon-gamma (IFN-γ) and lipopolysaccharide (LPS). Following a 48-hour activation period prior to infection, the monolayers were infected, and intracellular bacterial loads were determined 24 hours post-infection. Under these conditions, co-culture with activated THP-1 cells resulted in a pronounced and significant reduction in intracellular bacterial burden for both *L. monocytogenes* and *S.* Typhimurium compared with the corresponding epithelial-only controls (Figure 3C). Consistent with the reduced bacterial burden, LDH-based cytotoxicity measurements showed significantly lower infection-associated cytotoxicity in the presence of activated THP-1 cells for both pathogens (Figure 3D).

**Figure 3.**
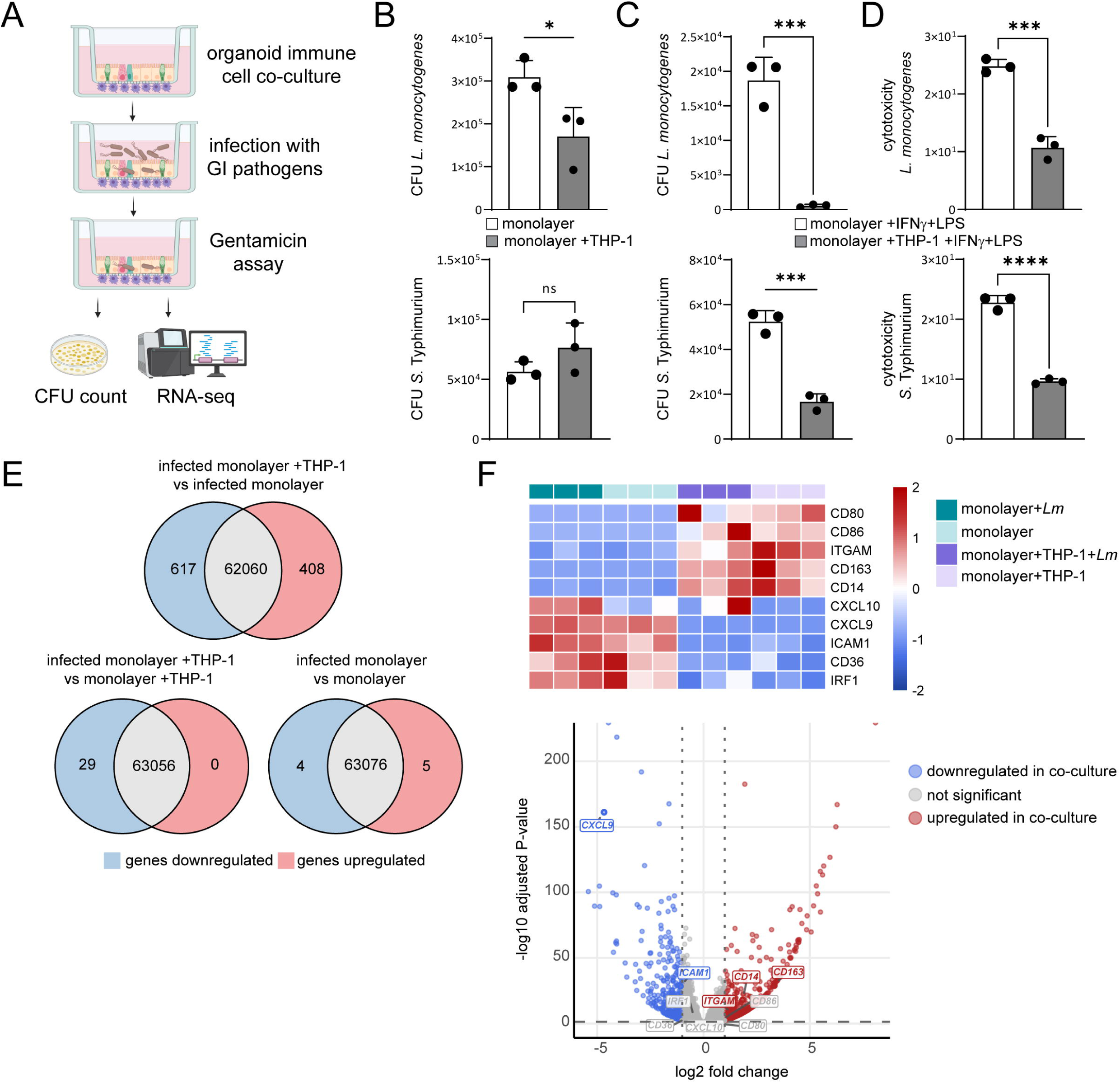
Activated macrophages restrict bacterial infection in intestinal organoid monolayers. (A) Schematic overview of the infection model. Differentiated organoid monolayers with or without basolateral THP-1 cells were infected apically, followed by gentamicin treatment and analysis of intracellular bacterial burden by CFU enumeration or transcriptomic profiling. (**B**) Intracellular *L. monocytogenes* (top) and *S.* Typhimurium (bottom) burden in organoid monolayers cultured with or without PMA-differentiated THP-1 cells at 24 h post-infection (MOI 10). Data are presented as mean ± SD. Two-tailed unpaired Student’s t-test. *L. monocytogenes*: representative of ≥3 independent experiments with three replicates per group; *S.* Typhimurium: one experiment with three replicates per group. (**C**) Intracellular *L. monocytogenes* (top) and *S.* Typhimurium (bottom) burden in organoid monolayers cultured with or without THP-1 cells following 48 h treatment with 20 ng/mL IFN-γ and 100 ng/mL LPS. Bacterial burden was determined 24 h post-infection (MOI 10). Data are presented as mean ± SD. Two-tailed unpaired Student’s t-test. Representative of ≥3 independent experiments with three replicates per group. (**D**) Infection-associated cytotoxicity of IFN-γ/LPS-treated organoid monolayers cultured with or without THP-1 cells following infection with *L. monocytogenes* (top) or *S.* Typhimurium (bottom), determined by LDH release at 24 h post-infection. Data are presented as mean ± SD. One-way ANOVA followed by Tukey’s multiple-comparisons test. Representative of two independent experiments with three replicates per group. Cytotoxicity shown as % of maximal LDH release (**E**) Diagrams summarizing differential gene expression for the indicated comparisons between infected and uninfected epithelial monocultures and THP-1 co-cultures. Genes with reduced expression are shown in blue, genes with increased expression in red, and genes without significant differential expression in grey. (**F**) Expression of selected macrophage-associated genes in epithelial monocultures and THP-1 co-cultures. Heatmap (top) shows row-wise Z-scores of normalized gene expression. Rows and columns were hierarchically clustered using Euclidean distance and Ward’s linkage method. The volcano plot (bottom) shows differential gene expression in infected co-cultures compared with infected epithelial monocultures. Selected macrophage-associated genes are labelled. Vertical dashed lines indicate log2 fold change > 1 and the horizontal dashed line indicates an adjusted p-value < 0.05. Genes meeting both criteria were considered significantly differentially expressed. n = 3 independent experiments per group.

To determine whether macrophage-associated restriction of bacterial burden could also be reproduced in a murine system, we additionally established a murine intestinal organoid monolayer co-culture with RAW264.7 macrophages (Figure S1A). Co-culture resulted in reduced Lgr5 expression, whereas expression of the epithelial lineage markers Alpi, Atoh1, and Muc2 remained unchanged, indicating only limited effects on the epithelial lineage-marker profile under these conditions (Figure S1B). Following prolonged IFN-γ/LPS activation, RAW264.7 co-culture significantly reduced intracellular S. Typhimurium burden compared with epithelial monolayers alone (Figure S1C). Fluorescence microscopy further revealed S. Typhimurium in both the epithelial and macrophage layers (Figure S1D), consistent with bacterial passage through the 1-µm Transwell membrane and access to the basolateral macrophage compartment.

### Transcriptional profiling of epithelial-macrophage co-cultures

To investigate the molecular changes associated with the reduced intracellular bacterial burden, we performed bulk RNA sequencing of human epithelial-macrophage co-cultures. Principal component analysis revealed that the presence of THP-1 cells was the dominant source of transcriptional variance between samples (Figure S2A). Given the close spatial association of epithelial cells and macrophages within the co-culture system, and the resulting contribution of transcripts from both cellular compartments to the bulk RNA pool, this separation was expected. In contrast, comparison of infected samples with their respective uninfected controls revealed only limited transcriptional changes in both epithelial monocultures and co-cultures (Figure 3E and S2A). Functional enrichment analysis of genes differentially expressed between co-cultures and epithelial monocultures identified several pathways associated with immune and myeloid cell functions, including pathways related to antigen processing and presentation, hematopoietic cell differentiation, phagocytosis, leukocyte migration, and osteoclast differentiation (Figure S2B). The enrichment of these pathways was consistent with the contribution of macrophage-derived transcripts to the bulk RNA-seq profiles of the co-culture samples. To further characterize the contribution of the macrophage compartment to the bulk RNA-seq profiles, we examined the expression of macrophage-associated genes (Figure 3F). As expected, several myeloid-enriched genes, including *CD14*, *CD163*, and *ITGAM*, showed increased expression in the co-culture samples, consistent with the presence of macrophage-derived transcripts. In contrast, several genes expressed by both epithelial and myeloid cells, including *CXCL9*, *CXCL10*, *ICAM1*, *IRF1*, and *CD36*, showed reduced expression in the co-cultures. Importantly, macrophage-associated and immune-response genes therefore displayed heterogeneous rather than uniformly increased expression patterns, indicating that the transcriptional differences between epithelial monocultures and co-cultures cannot be explained solely by the contribution of an additional cellular compartment. Instead, the co-culture was characterized by a distinct composite transcriptional profile consistent with altered gene regulation in the presence of macrophages.

### Co-culture is associated with coordinated changes in inflammatory and antimicrobial gene programs

Within the distinct transcriptional signature of the co-culture, one of the most prominent changes was a coordinated reduction in the expression of HLA-D genes associated with MHC class II-mediated antigen presentation (Figure 4A). This included several components of the MHC class II machinery, which showed consistently lower expression in co-cultures compared with epithelial monocultures. Given that intestinal epithelial cells can express MHC class II molecules in response to inflammatory signals (20; 21), the coordinated reduction of HLA-D gene expression may indicate that the presence of macrophages modulates the inflammatory activation state within the co-culture. We additionally observed prominent changes in the expression of *S100A8* and *S100A9*, which encode the two subunits of the antimicrobial complex calprotectin (Figure 4B). Calprotectin has both inflammatory and antimicrobial functions and can restrict bacterial growth, including *L. monocytogenes* and *S.* Typhimurium (22; 23). The differential expression of *S100A8* and *S100A9* therefore further suggests that macrophage co-culture alters inflammatory and antimicrobial programs within the system. Whether these changes contribute directly to the reduced intracellular bacterial burden observed in the co-culture remains to be determined.

**Figure 4.**
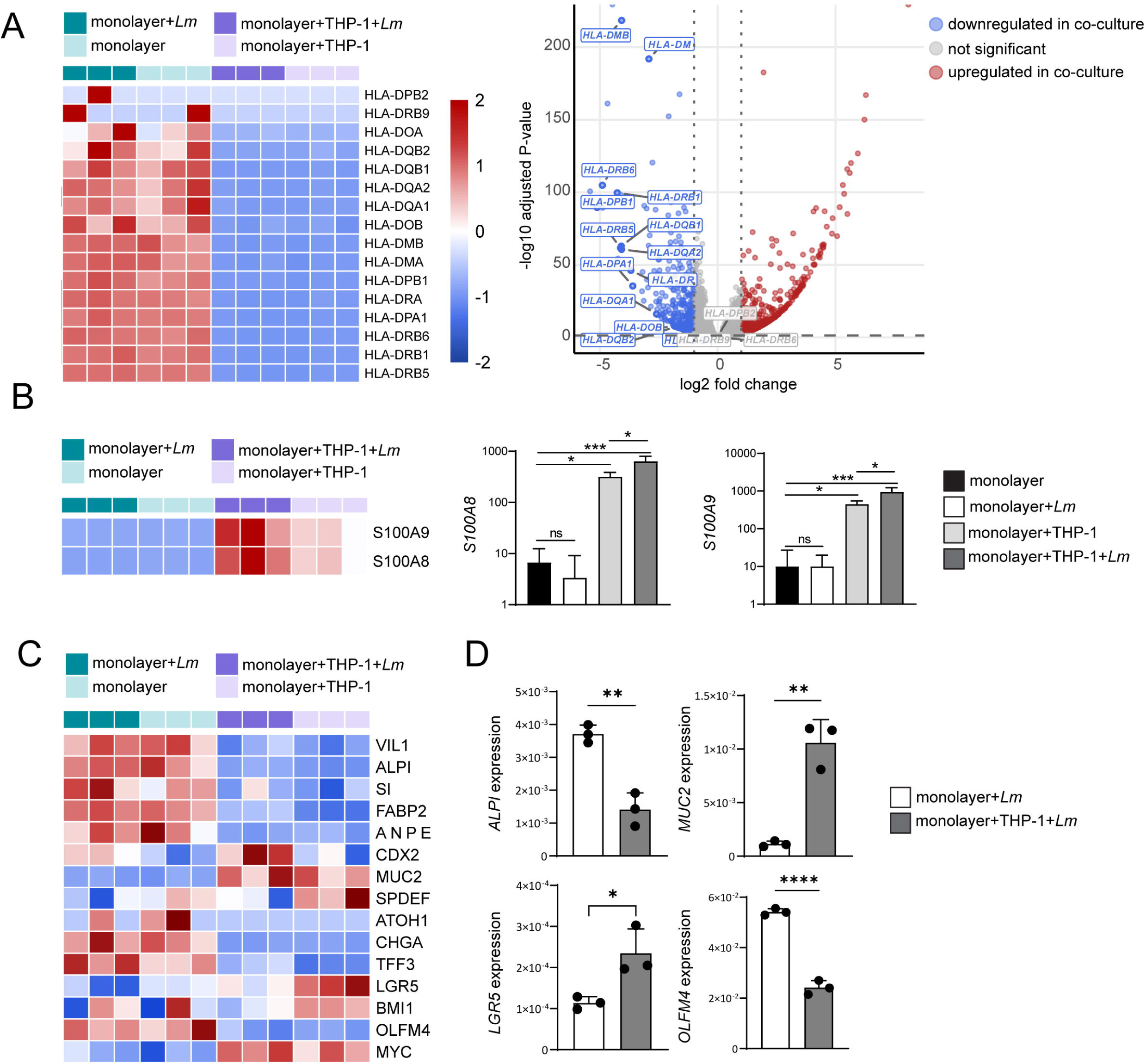
Macrophage co-culture under inflammatory conditions is associated with coordinated transcriptional changes. (**A**) Expression of HLA-D/MHC class II-associated genes in epithelial monocultures and THP-1 co-cultures under uninfected and *L. monocytogenes*-infected conditions. The heatmap (left) shows row-wise Z-scores of normalized gene expression; rows and columns were hierarchically clustered using Euclidean distance and Ward’s linkage method. The volcano plot (right) shows differential gene expression in infected co-cultures compared with infected epithelial monocultures. Selected HLA-D genes are labelled. Vertical dashed lines indicate log2 fold change > 1 and the horizontal dashed line indicates an adjusted p-value < 0.05. Genes meeting both criteria were considered significantly differentially expressed. n = 3 independent experiments per group. (**B**) Expression of *S100A8* and *S100A9* under the same experimental conditions. The heatmap (left) shows row-wise Z-scores of normalized gene expression. Bar graphs (right) show DESeq2 size-factor-normalized counts for *S100A8* and *S100A9* under the indicated conditions. Data are presented as mean ± SD. One-way ANOVA followed by Tukey’s multiple-comparisons test. n = 3 independent replicates per group. (**C**) Expression of selected epithelial lineage- and state-associated genes under the same experimental conditions. The heatmap shows row-wise Z-scores of normalized gene expression. Rows and columns were hierarchically clustered using Euclidean distance and Ward’s linkage method. n = 3 biological replicates per group. (**D**) Relative expression (2^-ΔCt^) of *ALPI*, *MUC2*, *LGR5*, and *OLFM4* in IFN-γ/LPS-treated, L. monocytogenes-infected human colon organoid monolayers cultured with or without THP-1 cells, determined by RT-qPCR. Data are presented as mean ± SD. Two-tailed unpaired Student’s t-test. One experiment with three replicates per group.

### Macrophage co-culture under inflammatory activation is associated with changes in epithelial lineage-associated gene expression

Whereas macrophage co-culture under basal conditions resulted in only limited changes in epithelial lineage markers (Figure 2C), a more pronounced shift was observed under the IFN-γ/LPS-activated infection conditions used for the transcriptomic analyses. Differential expression of several epithelial lineage- and state-associated genes, including *ALPI*, *ATOH1*, *MUC2*, *CHGA*, *LGR5*, *OLFM4*, and *MYC*, indicated a broader alteration of the epithelial transcriptional profile in the presence of macrophages (Figure 4C). To validate this shift using an independent method, representative epithelial markers were analyzed by RT-qPCR (Figure 4D). Consistent with the RNA-seq data, *ALPI* expression was reduced, whereas *MUC2* expression was increased in the co-cultures, supporting a shift from an enterocyte-associated toward a more goblet cell-associated transcriptional signature. Interestingly, the stem/progenitor-associated markers LGR5 and OLFM4 showed opposing expression patterns, with increased *LGR5* and reduced *OLFM4* expression in the co-culture, which was likewise confirmed by RT-qPCR. This divergence suggests that the observed changes do not simply reflect an expansion or loss of the conventional intestinal stem cell compartment, but may instead indicate an altered stem/progenitor-associated state under these conditions.

Together, these findings show that activated macrophage co-culture is associated with reduced intracellular bacterial burden and cytotoxicity, accompanied by coordinated changes in inflammatory, antimicrobial, and epithelial lineage-associated transcriptional programs.

## Discussion

In this study, we established a human intestinal organoid-derived monolayer–macrophage co-culture system that combines a polarized colonic epithelial barrier with basolaterally positioned macrophage-like cells and allows controlled apical infection with invasive bacterial pathogens. Under basal conditions, incorporation of THP-1 cells had only minor effects on epithelial lineage-associated gene expression and did not impair barrier integrity. In contrast, under inflammatory conditions, activated macrophages significantly reduced the intracellular burden of both *L. monocytogenes* and *S.* Typhimurium and this was accompanied by reduced infection-associated cytotoxicity. Consistent with the human system, activated RAW264.7 macrophages also reduced *S*. Typhimurium burden in murine intestinal organoid monolayers, supporting the transferability of the co-culture approach across species. Moreover, visualization of bacteria in both the epithelial and macrophage compartments suggests that pathogen control in this system may involve direct macrophage–pathogen interactions in addition to macrophage-mediated modulation of epithelial responses. Transcriptional profiling further demonstrated that macrophage co-culture was associated with coordinated changes in inflammatory, antimicrobial, and epithelial lineage-associated gene programs.

Several intestinal epithelial-immune cell co-culture systems have previously been developed. Noel et al. established a human intestinal stem cell-derived enteroid monolayer co-culture with primary monocyte-derived macrophages and demonstrated its applicability to intestinal physiology and bacterial host–pathogen interactions (18). This approach was subsequently further developed as a reproducible Transwell-based system incorporating macrophages or neutrophils beneath human enteroid or colonoid monolayers (19), and related models have been used to investigate coordinated epithelial–neutrophil responses to *Shigella* infection (17). In a different approach, Barrila et al. developed a three-dimensional HT-29/U937 co-culture model and observed reduced *Salmonella* colonization in the presence of macrophages (24). More recent macrophage-organoid co-culture models have further demonstrated that macrophages can influence epithelial homeostasis, inflammation, and regenerative responses (25; 26), and a recent murine small-intestinal organoid model incorporated macrophage co-culture to investigate epithelial and immune responses during *Mycobacterium avium* infection (27). Our model extends these approaches by combining an adult stem cell-derived human colonic monolayer with macrophage-like cells positioned directly at the basolateral side of the epithelial barrier and applying controlled infections with *L. monocytogenes* and *S*. Typhimurium together with functional and transcriptomic readouts.

A prominent transcriptional feature of the activated co-culture was the coordinated reduction of HLA-D genes involved in MHC class II antigen presentation. Intestinal epithelial cells can express MHC class II molecules, particularly in response to inflammatory signals, and epithelial MHC class II expression has been shown to influence intestinal immune responses and the severity of experimental inflammatory and infectious colitis (21). Thus, the reduced HLA-D signature observed in our co-culture may reflect a macrophage-associated modulation of the inflammatory activation state rather than a simple amplification of inflammatory signaling. This interpretation remains hypothetical, but suggests that the presence of macrophages may alter how the epithelial-immune system responds to a strong inflammatory environment. In addition, *S100A8* and *S100A9*, encoding the calprotectin complex, were prominently altered in the co-culture. Calprotectin has well-established antimicrobial functions and has been implicated in host responses to both *Listeria* and *Salmonella* infections (22; 23). While these changes are therefore compatible with modulation of an antimicrobial program, the present data do not establish a causal contribution of calprotectin to the reduced bacterial burden.

The presence of activated macrophages was also associated with changes in epithelial lineage-associated gene expression. Reduced *ALPI* and increased *MUC2* expression suggest a shift from an enterocyte-associated toward a more goblet cell-associated transcriptional signature. In parallel, *LGR5* and *OLFM4* showed opposing regulation, arguing against a simple expansion or depletion of a conventional intestinal stem cell compartment and potentially indicating an altered stem/progenitor-associated state. Immune-derived signals are known to influence intestinal stem cell renewal and differentiation (28), and recent work has identified macrophages as regulators of epithelial regeneration following intestinal injury (25). It is therefore conceivable that macrophage-derived signals contribute to the epithelial transcriptional changes observed here, particularly under inflammatory conditions. However, defining the cellular and molecular mechanisms underlying this shift will require additional studies.

Several limitations should be considered when interpreting these findings. First, the bulk RNA-seq profiles of the co-culture contain transcripts derived from both epithelial and macrophage compartments. Consequently, transcriptional changes cannot be unequivocally assigned to a specific cell type, and altered gene expression may reflect both changes within individual cell populations and changes in their relative contribution to the RNA pool. Similarly, the lineage-associated changes detected by RT-qPCR cannot distinguish altered gene expression within epithelial cells from changes in epithelial cellular composition. Future single-cell or spatial approaches could resolve these effects more directly. Second, THP-1-derived macrophage-like cells provide a reproducible experimental system but do not fully reproduce the phenotype and functional diversity of primary tissue-resident intestinal macrophages. Finally, the pronounced epithelial changes were observed under IFN-γ/LPS-activated infection conditions and should therefore be interpreted as context-dependent rather than as a general effect of macrophage co-culture. Extension of the system to primary or patient-matched macrophages and additional organoid donors will be important to determine the generalizability of these observations.

Despite these limitations, the model provides a modular platform in which the epithelial barrier, immune cell compartment, and luminal pathogen exposure can be experimentally controlled. Such systems may be adapted to different pathogens, immune cell populations, inflammatory conditions, and patient-derived organoids and could therefore complement animal models in mechanistic studies of intestinal infection and inflammation. In accordance with the 3R principles, organoid-based co-culture approaches may help reduce the number of animal experiments required for mechanistic screening while providing a human-relevant system for selected experimental questions (29). Further incorporation of primary immune cells, patient-derived material, and cell type-resolved molecular analyses should expand the translational potential of this approach. Overall, our findings demonstrate that intestinal organoid–macrophage co-cultures can reproduce functional epithelial–immune interactions during bacterial infection and provide a tractable platform for investigating how immune cells shape intestinal barrier responses to pathogenic challenge.

## Acknowledgments

Sequencing data used in this publication were generated by the Genomics & Single-Cell Core Unit (GENSEC) at Hannover Medical School. Lisa F. Goertz was supported by the Hannover Biomedical Research School (HBRS) and the Center for Infection Biology (ZIB). This work was funded by zukunft.niedersachsen (Federal State of Lower Saxony), R2N.Micro-Replace-Systems.

## Author contributions: CRediT

LFG: Investigation, Methodology, Formal analysis, Visualization, Writing – original draft; MC: Investigation; MB: Methodology, Resources; GAG: Methodology, Resources; ML: Conceptualization, Funding acquisition, Methodology, Supervision, Visualization, Writing – review and editing.

## Declaration of Generative AI and AI-assisted technologies in the writing process

During the preparation of this work the authors used ChatGpt (OpenAI) under full human supervision for proofreading, editing and reformatting of the final manuscript. After using this tool, the authors reviewed and edited the content as needed and take full responsibility for the content of the publication

## Methods

### Human and murine colon organoid culture and monolayer differentiation

Human colon biopsies were obtained from a healthy donor. Written informed consent was obtained in accordance with the approval of the ethics committee of Hannover Medical School (MHH; permit no. 3082-2016). Murine colonic organoids were generated from C57BL/6J mice bred and maintained under specific pathogen-free conditions at the Central Animal Facility of Hannover Medical School. Animals were housed and handled in accordance with the German Animal Welfare Act and Directive 2010/63/EU. Animals were sacrificed exclusively for scientific organ collection in accordance with §4(3) TierSchG and §2 TierSchVersV (notification no. 2023/244). Human colonic organoids and organoid-derived monolayers were cultured as described previously (30). Briefly, three-dimensional organoids were passaged every seven days. Organoids were dissociated into single cells using 0.05% trypsin/EDTA (Sigma-Aldrich) in combination with mechanical dissociation. A total of 1.5 × 10^4^ cells in 24 µL of organoid medium were mixed with 56 µL Matrigel (Corning) and seeded into pre-warmed 24-well plates. After polymerization for 30 min at 37 °C, pre-warmed organoid medium was added and replaced every other day. For establishment of murine organoids, colonic tissue was cut into small pieces and crypts were isolated using freshly prepared crypt-chelating buffer containing 10 mM EDTA in DPBS. Murine organoids were routinely passaged by mechanical dissociation and reseeded at a ratio of 1:3 to 1:4. Seven days before monolayer generation, murine organoids were passaged using the same dissociation procedure as described above.

For generation of organoid-derived monolayers, organoids were dissociated into single cells as described above. Transwell inserts with a pore size of 1 µm (Merck/Millipore) were coated with Matrigel diluted 1:25 in PBS for 3-24 h at 37 °C. Immediately before seeding, the remaining coating solution was removed. A total of 1.5 × 10^5^ cells in 200 µL monolayer medium were seeded into the apical compartment, and 800 µL medium was added to the basolateral compartment. Medium was replaced every 2-3 days. After six days, confluent monolayers were switched to differentiation medium and cultured for an additional 48 h before experimental use. Murine organoid-derived monolayers were generated and cultured using the same procedure. Organoid culture, monolayer, and differentiation media were prepared as described previously (30). Detailed medium compositions are provided in Supplementary Table S1.

### Organoid–macrophage co-culture and macrophage activation

Confluent organoid-derived monolayers were used for co-culture six days after seeding. THP-1 cells were differentiated into macrophage-like cells by treatment with 500 ng/mL phorbol 12-myristate 13-acetate (PMA; Sigma-Aldrich) for 48-72 h prior to establishment of the co-culture. PMA-differentiated THP-1 cells were collected, and 7 × 10^4^ cells were resuspended in 30 µL monolayer medium. Medium was removed from the Transwell inserts, which were subsequently inverted and placed in a 12-well plate. 30 µL of medium containing THP-1 cells were applied directly to the basolateral surface of the Transwell membrane. Control inserts received an equivalent volume of medium without immune cells. Following 4 h of incubation to allow cell attachment, residual medium was removed, the inserts were returned to their original orientation, and 200 µL and 800 µL differentiation medium were added to the apical and basolateral compartments, respectively. Co-cultures were maintained for 48 h before experimental use. For macrophage activation experiments, 100 ng/mL lipopolysaccharide (LPS; Sigma-Aldrich) and 20 ng/mL IFN-γ (PeproTech) were added to the basolateral differentiation medium during the 48-h co-culture period. Corresponding epithelial monoculture controls received the same concentrations of IFN-γ and LPS. For murine co-cultures, RAW264.7 macrophages were used without PMA differentiation. RAW264.7 cells were pre-activated with 100 ng/mL LPS and 10 ng/mL IFN-γ for 15-16 days prior to co-culture establishment. A total of 3 × 10^4^ pre-activated RAW264.7 cells were applied to the basolateral side of murine organoid-derived monolayers using the same co-culture procedure as described above. During the subsequent 48-h co-culture period, 100 ng/mL LPS and 10 ng/mL IFN-γ were maintained in the basolateral differentiation medium. Corresponding epithelial monoculture controls received the same treatment. Epithelial barrier integrity was monitored by transepithelial electrical resistance (TEER) measurements using a volt– ohm meter with electrodes positioned in the apical and basolateral compartments. TEER values were corrected for the surface area of the Transwell membrane.

### Bacterial infection and gentamicin protection assay

Overnight cultures of *S.* Typhimurium strain SL1344 were diluted 1:30 in Luria–Bertani (LB) broth (Sigma-Aldrich) and grown to an optical density at 600 nm (OD600) of 1.0. Overnight cultures of *L. monocytogenes* EGD were diluted 1:10 in brain heart infusion (BHI) medium (Thermo Fisher Scientific) and grown to an OD600 of 0.5. One hour before infection, the medium in the Transwell inserts was replaced with infection medium. Immediately before infection, the medium was removed from the apical compartment and replaced with 100 µL infection medium containing bacteria at a multiplicity of infection (MOI) of 10. For *L. monocytogenes* infection, Transwell inserts were centrifuged at 300 × g for 5 min at room temperature using reduced acceleration and braking. Following incubation for 1 h at 37 °C and 5% CO2, the inserts were washed with PBS and fresh infection medium containing 30 µg/mL gentamicin (Merck) was added to both the apical and basolateral compartments. For *S.* Typhimurium infection, monolayers were incubated with bacteria for 30 min before washing with PBS. Fresh infection medium containing 100 µg/mL gentamicin was subsequently added to both compartments. After an additional 1 h, the medium was replaced with infection medium containing 10 µg/mL gentamicin. Murine organoid-RAW264.7 co-cultures were infected with *S*. Typhimurium strain SL1344 using the same infection and gentamicin protection protocol as described above. At 24 h post-infection, Transwell inserts were washed with PBS and cells were lysed in distilled water containing 0.1% Triton X-100 (Sigma-Aldrich) and 0.05% sodium dodecyl sulfate (AppliChem). Lysates were serially diluted in PBS and plated on BHI agar for *L. monocytogenes* or LB agar for S. Typhimurium. Following incubation, intracellular bacterial loads were determined by enumeration of colony-forming units (CFUs).

### RNA isolation and cDNA synthesis

Total RNA was isolated using the RNeasy Mini Kit (Qiagen) according to the manufacturer’s instructions. For cDNA synthesis, RNA was reverse-transcribed using SuperScript™ IV Reverse Transcriptase (Invitrogen/Life Technologies). RNA was combined with random primers (Thermo Fisher Scientific) and dNTPs (Carl Roth) and incubated at 65 °C for 5 min, followed by cooling on ice. Subsequently, 5× SuperScript IV buffer, DTT, RNaseOUT™ Recombinant Ribonuclease Inhibitor, and SuperScript™ IV Reverse Transcriptase were added. Reverse transcription was performed by sequential incubation at 23 °C, 55 °C, and 80 °C.

### Quantitative real-time PCR

Quantitative real-time PCR (qPCR) was performed using iQ™ SYBR® Green Supermix (Bio-Rad) on a CFX Opus 96 Real-Time PCR System (Bio-Rad). Gene expression was determined using gene-specific primers. Relative expression levels were calculated using the 2^-ΔCt^-method and normalized to GAPDH for human samples and Actb for murine samples. Primer sequences are provided in Supplementary Table S2.

### RNA sequencing and bioinformatic analysis

RNA sequencing was performed by the Research Core Unit Genomics at Hannover Medical School. For each sample, 250 ng of total RNA was used for poly(A) mRNA enrichment using the NEBNext Poly(A) mRNA Magnetic Isolation Module (New England Biolabs). Stranded cDNA libraries were subsequently generated using the NEBNext Ultra II Directional RNA Library Prep Kit for Illumina (New England Biolabs) according to the manufacturer’s instructions, with reaction volumes reduced to two-thirds of the recommended volumes. Libraries were barcoded using NEBNext Multiplex Oligos for Illumina Unique Dual Index Primer Pairs and amplified using nine cycles of PCR. Following library preparation, an additional purification step was performed using 1.2 x Agencourt AMPure XP beads (Beckman Coulter). Library fragment size distributions were assessed using the Bioanalyzer High Sensitivity DNA Assay (Agilent Technologies), and library concentrations were determined using the Qubit® dsDNA High Sensitivity Assay Kit (Thermo Fisher Scientific). Individually barcoded libraries were pooled, denatured, and sequenced on an Illumina NovaSeq X Plus platform using 2 x 61-bp paired-end sequencing. Sequencing yielded approximately 25-35 million read pairs per sample. BCL files were converted to FASTQ format using bcl2fastq Conversion Software v2.20.0.422 (Illumina). Raw sequencing data were processed using the nf-core/rnaseq pipeline (version 3.9), which performs read preprocessing, alignment, quantification, and quality control. Reads were aligned to the human reference genome GRCh38.p14 using genome annotation release 46 obtained from GENCODE. Principal component analysis (PCA) was performed as part of the nf-core/rnaseq workflow using DESeq2 and was used to visualize transcriptional variation between samples.

Normalization and differential gene expression analysis were performed using DESeq2 (Galaxy tool version 2.11.40.6) on the internal Galaxy platform of the Research Core Unit Genomics, Hannover Medical School. DESeq2 was run using default settings, except that normalized count tables were generated and outlier replacement, outlier filtering, and independent filtering were disabled. Experimental conditions were included as levels of the primary factor and pairwise comparisons were performed between the respective groups. Unless otherwise indicated, genes were considered differentially expressed based on an adjusted p-value < 0.05 and the fold-change thresholds specified for the respective analyses.

### RNA-seq data visualization and pathway analysis

Downstream analysis and visualization of differential gene expression data were performed in R. Volcano plots were generated using the R packages ggplot2, ggrepel, and dplyr. Heatmaps were generated using pheatmap, with data handling performed using dplyr. For heatmap visualization, expression values were transformed to row-wise Z-scores, and hierarchical clustering of genes and samples was performed using Euclidean distance and Ward’s linkage method. Venn diagrams illustrating overlaps between differentially expressed gene sets were generated using ggplot2 and ggforce. Functional enrichment analysis of differentially expressed genes was performed using the clusterProfiler package together with the human genome annotation package org.Hs.eg.db. Kyoto Encyclopedia of Genes and Genomes (KEGG) pathway enrichment was analyzed separately for upregulated and downregulated genes. Pathways with an adjusted p-value < 0.05 were considered significantly enriched. Graphical representations were generated using ggplot2 and exported using svglite.

### THP-1 and RAW264.7 cell culture

THP-1 cells were maintained in RPMI 1640 medium (Thermo Fisher Scientific) supplemented with 10% fetal calf serum (FCS; Bio&SELL GmbH) and 55 µM 2-mercaptoethanol (Life Technologies). Cells were maintained in suspension culture and diluted approximately once per week by removing 90% of the cell suspension and replacing it with fresh culture medium. RAW264.7 cells were maintained in DMEM (Thermo Fisher Scientific) supplemented with 10% FCS. Cells were passaged twice weekly by reseeding approximately 10% of the cells into a new culture flask and adding fresh medium.

### Immunofluorescence staining

Prior to establishment of the co-culture, THP-1 cells were fluorescently labelled with carboxyfluorescein succinimidyl ester (CFSE; Thermo Fisher Scientific). Cells were collected and resuspended in 1 mL PBS containing CFSE at a concentration of 5 µM. Following incubation for 7 min at 37 °C, cells were placed on ice for 5 min. Subsequently, 5 mL of cell culture medium was added and cells were incubated for an additional 5 min before centrifugation and resuspension in fresh culture medium. For murine co-cultures, RAW264.7 macrophages were labelled with CellTrac Far Red (Thermo Fisher Scientific) immediately before co-culture establishment. Cells were collected by centrifugation at 300 x g for 7 min at room temperature, resuspended in 1 mL PBS containing 1 µM CellTrace Far Red, and incubated for 20 min at 37 °C protected from light. Subsequently, 5 mL cell culture medium was added, cells were incubated for an additional 5 min, centrifuged, and resuspended in fresh culture medium. For fluorescence imaging of bacterial localization, a GFP-expressing *S*. Typhimurium NCTC 12023 (14028S) strain was used.

For immunofluorescence staining of epithelial monolayers, cells were fixed with 4% paraformaldehyde (Thermo Fisher Scientific) in PBS for 15 min. Transwell membranes were excised from the inserts and permeabilized in staining buffer for 30 min. Membranes were subsequently incubated with a primary antibody against zonula occludens-1 (ZO-1; Thermo Fisher Scientific), diluted in staining buffer, for 1 h at room temperature. Following washing with PBS, membranes were incubated for 30 min with Alexa Fluor 633-conjugated goat anti-rabbit IgG secondary antibody together with DAPI and phalloidin (Thermo Fisher Scientific) and/or Ulex europaeus agglutinin I (UEA-I; Vector Laboratories), as indicated. For imaging of infected murine co-cultures, nuclei were stained with DAPI and F-actin with phalloidin. GFP-expressing *S*. Typhimurium and CellTrace Far Red-labelled RAW264.7 cells were detected directly. After a final wash with PBS, membranes were mounted on glass slides using mounting medium and covered with coverslips. Slides were analyzed on a ZEISS LSM 980 with Airyscan 2.

### LDH cytotoxicity assay

Infection-associated cytotoxicity was assessed using the CyQUANT™ LDH Cytotoxicity Assay Kit (Thermo Fisher Scientific). At the indicated time point, 50 µL of apical supernatant from infected organoid monolayers was transferred to a 96-well plate. Culture medium alone served as a background control, whereas supernatant from lysed uninfected organoid monolayers was used as a positive control for maximal LDH release. 50 µL of substrate solution was added to each well and incubated for approximately 20 min, until a strong colorimetric signal was observed in the positive control. The reaction was stopped by addition of 50 µL stop solution. Absorbance was measured at 490 nm and 680 nm using a Synergy HTX Multimode Reader (BioTek). Cytotoxicity was expressed relative to the maximal LDH release of the positive control, which was set to 100%, and calculated as (sample/positive control) x 100.

### Statistical analysis

Statistical analyses were performed using GraphPad Prism (GraphPad Software). Comparisons between two independent groups were performed using two-tailed unpaired Student’s t-tests. Multiple-group comparisons were performed using one-way analysis of variance (ANOVA) followed by Tukey’s multiple-comparisons test. Data are presented as mean ± SD unless otherwise indicated. Statistical significance was defined as follows: ns, p > 0.05; *, p ≤ 0.05; **, p ≤ 0.01; ***, p ≤ 0.001; and ****, p ≤ 0.0001. The statistical tests and numbers of independent experiments or biological replicates are specified in the respective figure legends.

### Data availability

The raw and processed RNA-seq data generated in this study have been deposited in the NCBI Gene Expression Omnibus (GEO) under accession number GSE345254.

## Supplemental information

Document S1. Figures S1-2, Tables S1-S2

