## Supplementary Figure S1-S2 and Tables S1-2 for "Activated macrophages restrict invasive bacterial infection in a human intestinal organoid co-culture model"

**Figures S1-S2**
**Tables S1-S2**

#### Supplemetary Figure 1

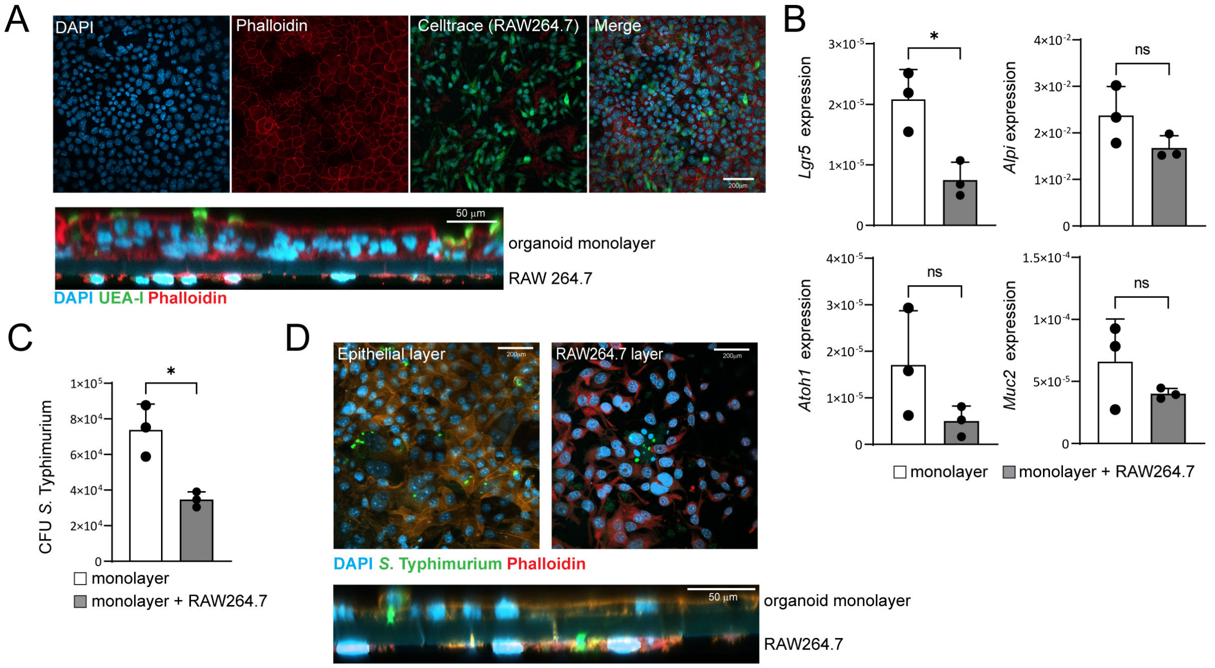


**Figure S1. Activated RAW264.7 macrophages restrict *S.* Typhimurium infection in murine intestinal organoid monolayers.** (**A**) Representative fluorescence images of murine intestinal organoid-derived monolayers co-cultured with RAW264.7 macrophages. Top view: nuclei (DAPI, blue), F-actin (phalloidin, red), and CellTrace-labelled RAW264.7 cells (green). Cross-section: nuclei (DAPI, blue), F-actin (phalloidin, red), and UEA-I (green), showing the epithelial monolayer and basolaterally positioned RAW264.7 macrophages. Scale bars as indicated. (**B**) Relative expression of *Lgr5*, *Alpi*, *Atoh1*, and *Muc2* in murine intestinal organoid monolayers cultured with or without RAW264.7 macrophages, determined by RT-qPCR. Data are presented as mean ± SD. Two-tailed unpaired Student's t-test. n = 3 independent experiments per group. (**C**) Intracellular *S.* Typhimurium burden in murine intestinal organoid monolayers cultured with or without RAW264.7 macrophages following prolonged IFN-γ/LPS activation. Bacterial burden was determined by CFU enumeration. Data are presented as mean ± SD. Two-tailed unpaired Student's t-test. n = 3 independent experiments per group. (**D**) Representative fluorescence images of *S.* Typhimurium-infected murine organoid-RAW264.7 co-cultures. Images show focal planes corresponding to the epithelial monolayer (left) and RAW264.7 macrophage layer (right), together with a cross-sectional Z-stack view (bottom). Nuclei are shown in blue, *S.* Typhimurium in green, and F-actin in red. Scale bars as indicated.

### Supplemetary Figure 2

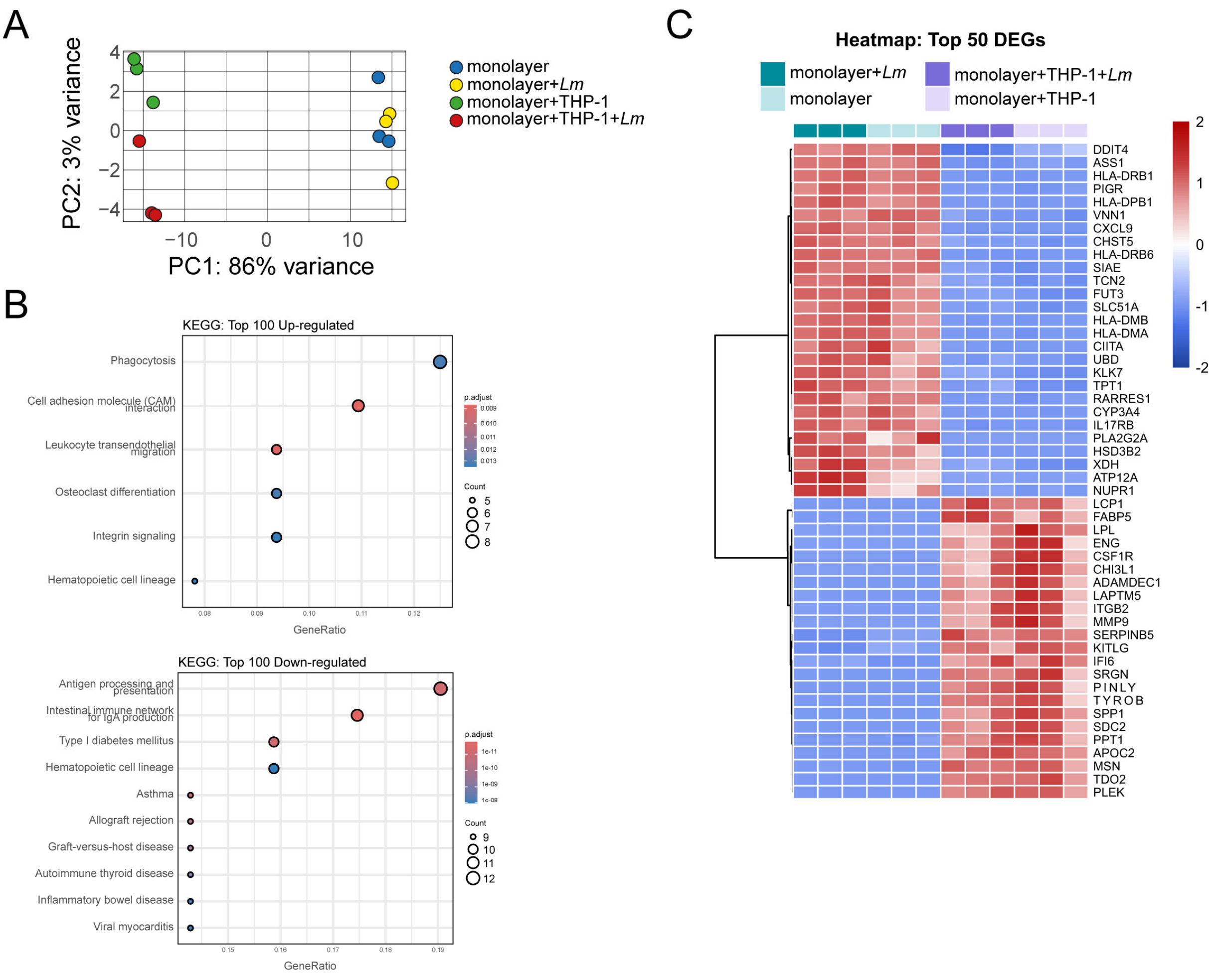


**Figure S2. Transcriptomic characterization of epithelial monocultures and THP-1 co-cultures.** (**A**) Principal component analysis (PCA) of RNA-seq samples from epithelial monocultures and THP-1 co-cultures under uninfected and *L. monocytogenes*-infected conditions. PCA was performed using DESeq2-normalized expression data. Each point represents one biological replicate. n = 3 independent experiments per group. (**B**) Functional enrichment analysis of differentially expressed genes in infected co-cultures compared with infected epithelial monocultures. The most significantly enriched KEGG pathways are shown separately for upregulated and downregulated genes. Enrichment analysis was performed using an over-representation approach, and pathways with an adjusted p-value < 0.05 were considered significantly enriched. n = 3 independent experiments per group. (**C**) Heatmap of the top 50 differentially expressed genes in co-cultures compared with epithelial monocultures. Expression levels are shown as row-wise *Z*-scores. Rows and columns were hierarchically clustered using Euclidean distance and Ward's linkage method. n = 3 independent experiments per group.

**Table S1. Media used for organoid culture**

Standard organoid medium

| Ingredient | Concentration |
|---|---|
| DMEM | 45% |
| L-WRN supernatant | 50 % |
| GlutaMax | 2 mM |
| HEPES | 10 mM |
| B-27 supplement | 1 x |
| N-acetyl-L-cysteine | 1 mM |
| EGF | 50 ng/mL |
| Y-27632 | 10 µM |
| A83-01* | 500 nM |
| SB202190* | 10 µM |
| Gastrin I* | 10 nM |

*A83-01, SB202190, and Gastrin I were included only in the human standard organoid medium.

Organoid monolayer medium

| Ingredient | Concentration |
|---|---|
| Advanced DMEM/F12 | 28% |
| L-WRN | 50 % |
| FCS | 20 % |
| GlutaMax | 2 mM |
| Y-27632 | 10 µM |
| EGF | 50 ng/mL |
| Streptomycin | 0.1 mg/mL |
| Penicillin | 100 U/mL |

Organoid monolayer differentiation medium

| Ingredient | Concentration |
|---|---|
| Advanced DMEM/F12 | 73% |
| L-WRN | 5 % |
| FCS | 20 % |
| GlutaMax | 2 mM |
| DAPT | 5 µM |
| EGF | 50 ng/mL |

**Table S2. Primers used for RT-qPCR**

| Gene | Forward primer | Reverse primer |
|---|---|---|
| ALPI (h) | GCAACCCTGCAACCCACCCAAGGAG | CCAGCATCCAGATGTCCCGGGAG |
| ATOH1 (h) | GCAATGTTATCCCGTCGTTCA | CCATCTGCAGGGTCTCATATTTG |
| GAPDH (h) | GGTCTCCTCTGACTTCAACA | AGCCAAATTCGTTGTCATAC |
| LGR5 (h) | AATCCCCTGCCCAGTCTC | CCCTTGGGAATGTATGTCAGA |
| MUC2 (h) | ATGCCCACCTCCTCAAAGAC | GTAGTTTCCGTTGGAACAGTGAA |
| OLFM4 (h) | TTCTCCTAGCCCTTCTGTTCTTCC | TTCCAAGCGTTCCACTCTGTCC |
| Actb (m) | TGTTACCAACTGGGACGACA | GGGGTGTTGAAGGTCTCAAA |
| Alpi (m) | AGGATCCATCTGTCCTTTGG | ACGTTGTATGTCTTGGACAG |
| Atoh1 (m) | GGGGTTGTAGTGGACGAGC | CGTTGTTGAAGGACGGGATAAC |
| Lgr5 (m) | GACAATGCTCTCACAGAC | GGAGTGGATTCTATTATTATGG |
| Muc2 (m) | GTCCGAAGTGTTACCCTGGA | CCAGGAGTGGAGAAGGTCAG |

(h) human, (m) murine
